# Open- and Closed-Loop Microcomplexes in Neocerebellum

**DOI:** 10.64898/2026.02.17.706406

**Authors:** Willem S. van Hoogstraten, Elías M. Fernandez Santoro, Lorena M. Hähner, Chris I. De Zeeuw

**Author notes:** These authors contributed equally to this work and share first authorship (Willem S. van Hoogstraten, Elías M. Fernandez Santoro).

## Abstract

Neocerebellum facilitates motor and cognitive behavior via olivocerebellar modules, in which Purkinje cells (PCs), cerebellar nuclei (CN) neurons and olivary cells are supposed to form exclusively closed-loops. Here, we provide evidence that parts of the modules may be organized in an open-loop feedforward fashion where PCs do not contact the CN neurons that provide feedback to the olive. The PCs in the open-loop microcomplexes exclusively target projection CN neurons, through which cerebellum modulates downstream brainstem regions controlling behavior. This novel cerebellar architecture, containing open– and closed-loop microcomplexes within single modules, offers a mechanism by which mossy fiber information can be differentially used either as feedforward control of behavior and/or as feedback to the olivary subnuclei that regulates activity and plasticity within the module involved. This potential for both integrated and segregated regulation of behavior and feedback within single modules enriches the computational repertoire of the olivocerebellar system.

## Introduction

The cerebellum is involved in coordination of many different types of behaviors, ranging from reflexive and voluntary movements, up to autonomic, emotional and cognitive efforts (Schmahmann 1996, Ito 2008, Fujita *et al*. 2020, van der Heijden *et al*. 2023). Classical tracing and electrophysiological studies of the past century have revealed that control of these different behaviors is mediated by specific olivocerebellar modules (Apps *et al*. 2018). The modules are formed by three-element loops, comprising a particular microzone of Purkinje cells (PCs), their target neurons in the cerebellar nuclei (CN), and a subnucleus in the inferior olive (IO) that provides the climbing fiber input to the PCs of the same microzone (Clendenin *et al*. 1974, Andersson and Oscarsson 1978, Apps and Garwicz 2005). Given that virtually all GABAergic CN projection neurons provide an inhibitory feedback to the IO subnuclei with a high level of topography (Ruigrok and Voogd 1990, Wang *et al*. 2023, van Hoogstraten *et al*. 2024), it has been assumed that the three-element PC-CN-IO loops within the modules are consistently organized in a closed-loop fashion (**Fig. 1**, left side). This implies that at the level of an individual microcomplex, which comprises the PCs and connected CN neurons of a particular micromodule (Apps and Garwicz 2005), the PCs are always supposed to contact both the excitatory neurons in the CN that project to particular premotor output areas (Teune *et al*. 2000) and the GABAergic CN neurons that provide feedback to an olivary subnucleus belonging to the same micromodule (Ruigrok and Voogd 1990, Wang *et al*. 2023, van Hoogstraten *et al*. 2024). Indeed, as the excitatory neurons in the CN that directly control behavior are spatially closely intermingled with the CN neurons that provide the olivary feedback (Uusisaari and De Schutter 2011), it is parsimonious to expect that the PCs that project to a particular part of the CN consistently innervate both types of neurons (De Zeeuw and Berrebi 1995, de Zeeuw and Berrebi 1996, Teune *et al*. 1998). However, this classical dogma on the architecture of the olivocerebellar modules would suggest that cerebellar control of behavior via a particular microcomplex always rides on top of an olivary loop that is closed, limiting its computational repertoire. Given the emerging evidence that neocerebellum can perform a great variety of complex computations (Zang and De Schutter 2023), we hypothesized that open-loop microcomplexes may exist inside their modules next to the closed-loop microcomplexes, allowing for segregated regulation of behavior and feedback within single modules and thereby enriching the computational repertoire (**Fig. 1**).

**Fig. 1.**
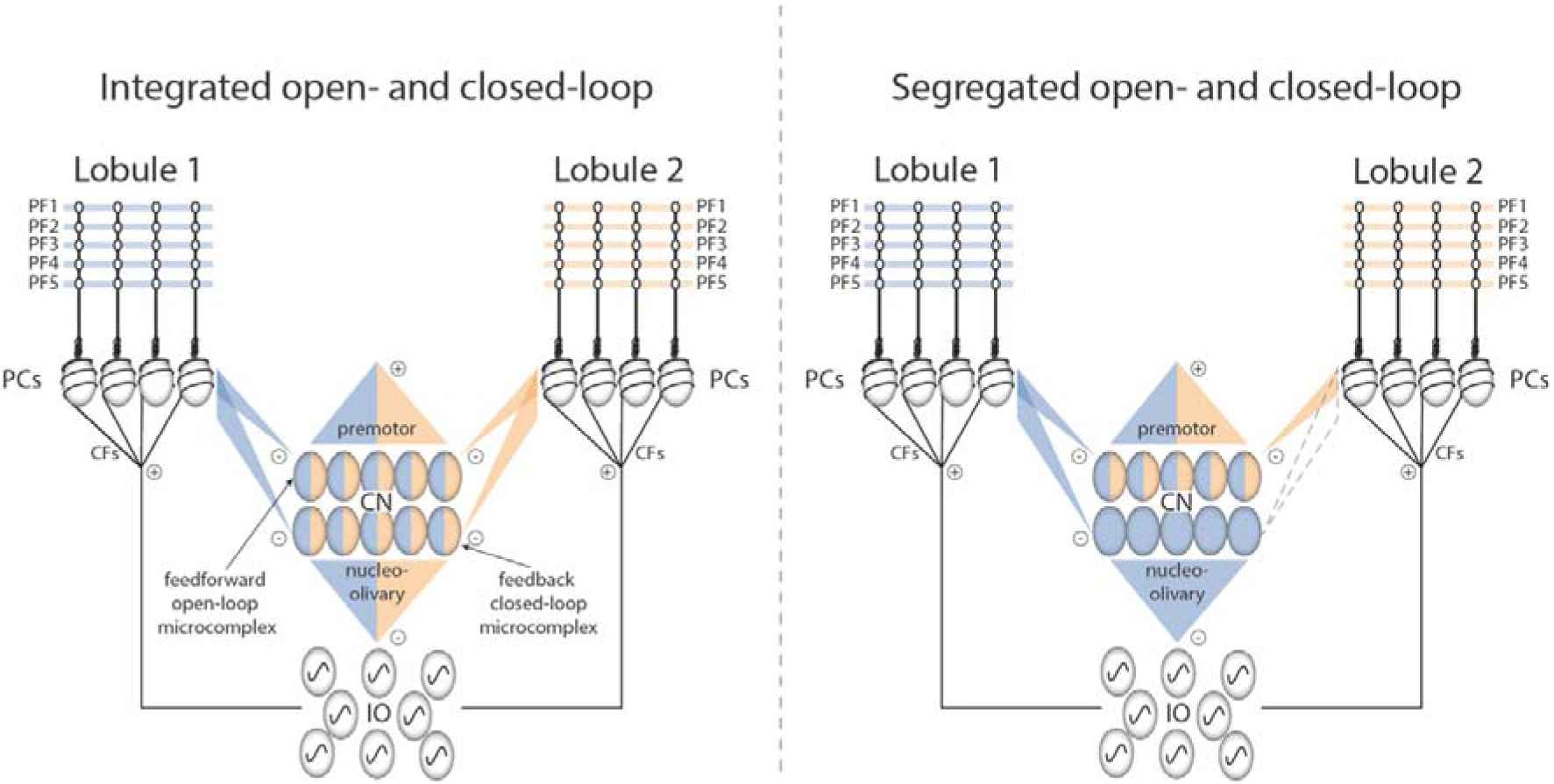
| Architecture of neocerebellum according to the classical dogma with open– and closed-loop microcomplexes that are always integrated (left) and according to our novel hypothesis with partially segregated open– and closed-loop microcomplexes (right). The traditional model on the left assumes that all Purkinje cells (PCs) of the cerebellar cortex (i.e., including all PCs of a module distributed across different lobules) provide feedback to the inferior olive (IO) via the GABAergic cerebellar nuclei (CN) cells and that all these PCs in addition provide feedforward projections to excitatory premotor output neurons in the CN. Our novel model on the right hypothesizes that some PCs of a module are involved in both the open– and closed-loop microcomplexes, but that other PCs of the same module are only involved in the open-loop microcomplex. The latter configuration would imply that the mossy fiber – parallel fiber (PF) inputs to a particular set of PCs in a particular lobule (either blue or yellow) determines what information is used to control the feedforward and/or feedback processing of a particular module and that regulation of the feedforward and feedback processing can be partially segregated.

## Results

We evaluated our hypothesis on segregated open– and closed-loop microcomplexes by doing transsynaptic tracing experiments. We made injections of the adeno-associated viral vector AAV1-CMV-HI-eGFP-Cre-WPRE-SV40 (30nL) into the neocerebellar cortex of adult, male and female, Ai14 reporter mice (n = 18), which are homozygous for the Rosa-CAG-LSL-tdTomato-WPRE conditional allele (Zingg *et al*. 2017, 2020, 2022). Unlike classical tracing experiments, this approach allows us to simultaneously trace the transsynaptically, anterogradely labeled somata and axons of CN cells and the retrogradely labeled cells in the IO of the same module (**Fig. 2a**). Large injections that contained histological damage at the injection site and that did not result in sufficient transsynaptic labeling of the superior cerebellar peduncle were excluded from further analyses (n = 3). For the analyses of the remaining 15 injections, we identified the microzonal location of each injection site based on the distribution of PC labeling in the neocerebellar cortex (**Fig. 2b**) as well as on the location of the retrogradely labeled cells in the IO (**Fig. 2c, d**) (Ruigrok 2003, Voogd *et al*. 2003, Owusu-Mensah *et al*. 2023). The delineation of the microzones was done with the use of anti-zebrin labeling (KCTD12) (Sugihara and Quy 2007, Fujita *et al*. 2014, Blot *et al*. 2023). All injections covered one or more parts of the paravermis and/or hemisphere of lobule simplex (n = 2; 4-B7 and 4-B8), crus 1 (n = 4; 302-B5, 236-A5, 302-B4 and A4), crus 2 (n = 3; 4-B5, 236-A6 and 4-B6) or the paramedian lobule (n = 5; 4-B10, 4-B9, 236-A3, B1, B2 and 395-B3), (for overview, see **Fig. 2e**). Likewise, most of the AAV injections we made were taken up by PCs from more than one microzone and each module was targeted by at least two injections (**Fig. 2e**): X-CX (n = 3; 236-A3, 4-B6 and 302-B4), C1 (n = 4; 236-A3, 295-B3, B2 and 4-B6), A2 (n = 12; 4-B7, 4-B8, A4, 302-B4, 236-A5, 302-B5, 236-A6, 4-B6, 236-A3, B1, B2 and 395-B3), C2 (n = 7; 4-B7, 236-A5, 302-B5, 302-B4, 236-A6, 236-A3 and B1), C3 (n = 4; 4-B7, 236-A6, 4-B9 and 236-A3), D1 (n = 5; 236-A5, 302-B5, 4-B5, 4-B9 and 236-A3), D0 (n = 2; 4-B5 and 4-B9), and D2 (n = 4; 302-B5, 4-B5, 4-B10 and 4-B9). In line with the localization of the injections in the cerebellar cortex, we observed that the labeling observed in the X-CX, A2 and C2 zones, C1 and C3 zones, and D zones corresponded with retrogradely labeled cells in the medial accessory olive (MAO), dorsal accessory olive (DAO) and principal olive (PO), respectively (**Fig. 2c, d, e**).

**Fig. 2.**
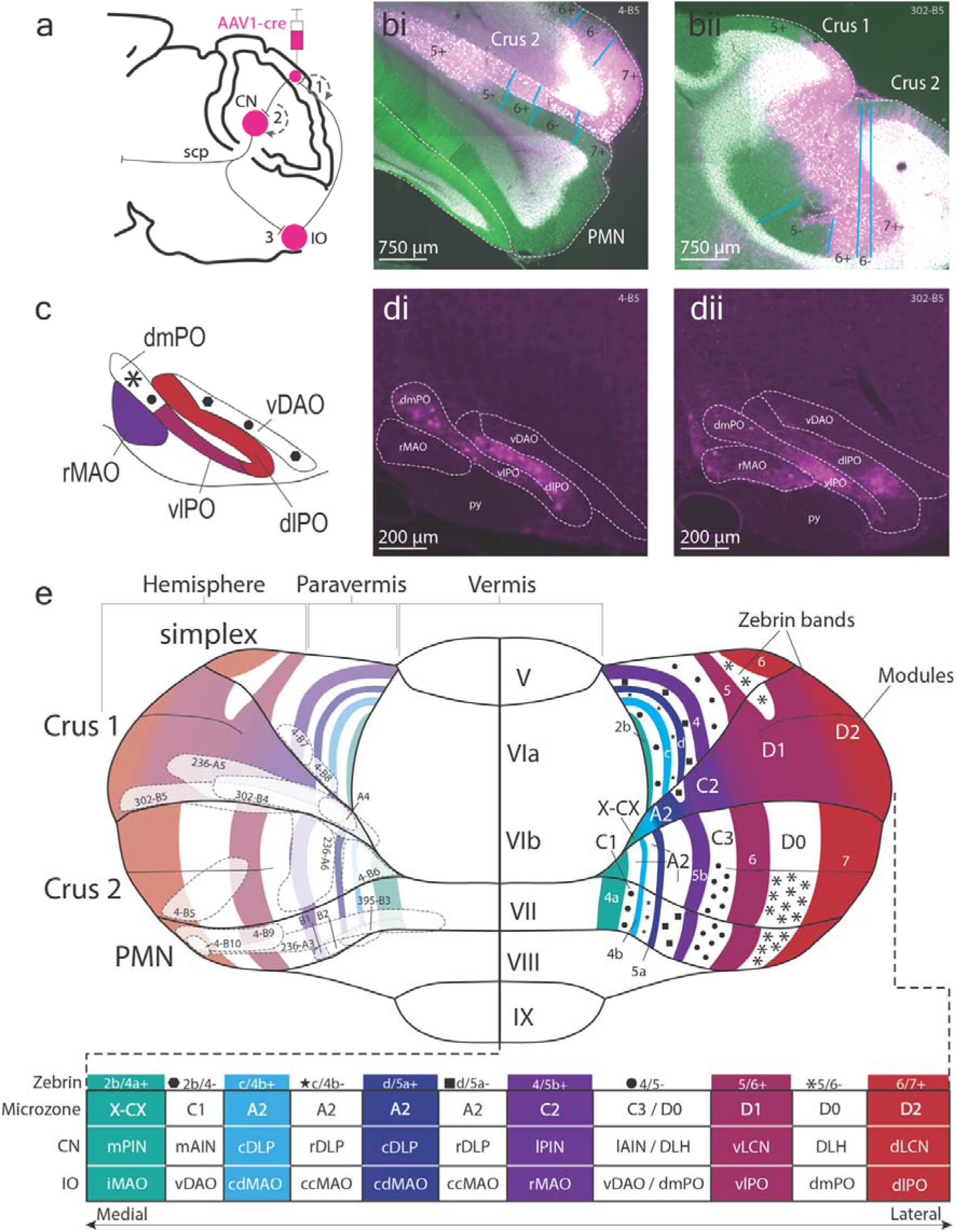
| Delineation and identification of injection sites with respect to modules and lobules. (**a**), Schematic representation of our pseudo-trisynaptic tracing approach. The first synapse is inferred from retrograde uptake by the climbing fibers in the cerebellar cortex, leading to retrograde transfection of cells in the inferior olive (IO). The second synapse is inferred from transsynaptic transfection of the cells in the cerebellar nuclei (CN) that receive input from the primarily transfected Purkinje cells (PCs). The third synapse is inferred from confocal imaging of anterogradely labeled varicosities in the IO, originating from the transsynaptically labelled CN cells, completing the pseudo-trisynaptic tracing trajectory. (**b**), Examples of injections (4-B5 in bi and 302-B5 in bii) in Crus 1 and 2 with green KCTD12 staining as reference frames to identify the zebrin zones. TdTomato is visualized in magenta, displaying the initial injection site. (**c**), Schematic drawing of a coronal view of the IO and its subnuclei; the colors and black markers highlight how the different olivary subnuclei provide the climbing fibers to specific microzones in the cerebellar cortex as indicated in panel e. (**d**), Example images of retrogradely labelled cells in the IO of injections 4-B5 (di) and 302-B5 (dii). (**e**), Schematic drawing of the cerebellar cortex (from dorsal perspective) where the zebrin bands and microzones (large white/black text) are represented together. The precise relations between the microzones, CN and IO subnuclei are visualized in different colors. This is summarized in the matrix at the bottom. For all injections, the delineations on a zebrin-map were confirmed based on the retrograde labeling in IO. Information for the approximate translation of zebrin-zones into modules was deduced from data obtained by us as well as others (Apps 1990, Buisseret-Delmas and Angaut 1993, Atkins and Apps 1997, Pardoe and Apps 2002, Voogd *et al*. 2003, Sugihara and Shinoda 2004, Voogd and Ruigrok 2004, Sarna *et al*. 2006, Sugihara and Quy 2007, Apps and Hawkes 2009, Sillitoe *et al*. 2012, Fujita *et al*. 2020, Owusu-Mensah *et al*. 2023, van Hoogstraten *et al*. 2024).

All injections resulted in transsynaptic anterograde labeling of cell-bodies in the CN (**Fig. 3a**) as well as of their connected axons in or adjacent to the superior cerebellar peduncle (**Fig. 3b**). The location and density of labeling within the superior cerebellar peduncle could be related to the location of the injection site as well as the location, shape and size of the transsynaptically labeled cell-bodies within the CN in that some injections, such as those of crus 1, resulted predominantly in dense labeling of the medial part of the superior cerebellar peduncle, whereas others, such as those of crus 2, resulted in relatively weak labeling of the more lateral part of the superior cerebellar peduncle (**Fig. 3b**). Since the inhibitory GABAergic nucleo-olivary fibers and the axons of the excitatory glutamatergic premotor neurons in the CN are predominantly located in the medial and lateral part of the superior cerebellar peduncle, respectively (Ruigrok and Voogd 1990, Teune *et al*. 2000, Apps and Garwicz 2005), these findings raise the possibility that open-loop microcomplexes indeed exist next to the typical closed-loop microcomplexes (**Fig. 1**). To further investigate this possibility, we next determined for each injection to what extent the distribution of the transsynaptic, anterograde labeling of GABAergic terminals in the IO corresponded to that of the retrograde labeling in the olive (**Fig. 3c, d**). If open-loop microcomplexes exist next to the closed-loop microcomplexes, one expects to find some injections that result in partially segregated anterograde and retrograde labeling in the olive, whereas others should result in a perfect overlap of both types of labeling. We did indeed find evidence for both types of distributions (**Table S1**). Nine Injections (236-A5, 302-B5, 302-B4, A4, 4-B9, 4-B10, 236-A3, B1 and B2) resulted in a perfect spatial overlap between the transsynaptic, anterograde labeling and the retrograde labeling in the IO (**Fig. 3cii**). The other six injections yielded a different pattern, with three injections (4-B8, 4-B9, and 4-B5) showing only partial overlap and the other three (4-B6, 236-A6, 395-B3) showing exclusively retrograde labeling in the IO, i.e., they did not show any transsynaptic, anterograde labeling at all (**Fig. 3ci**).

**Fig. 3.**
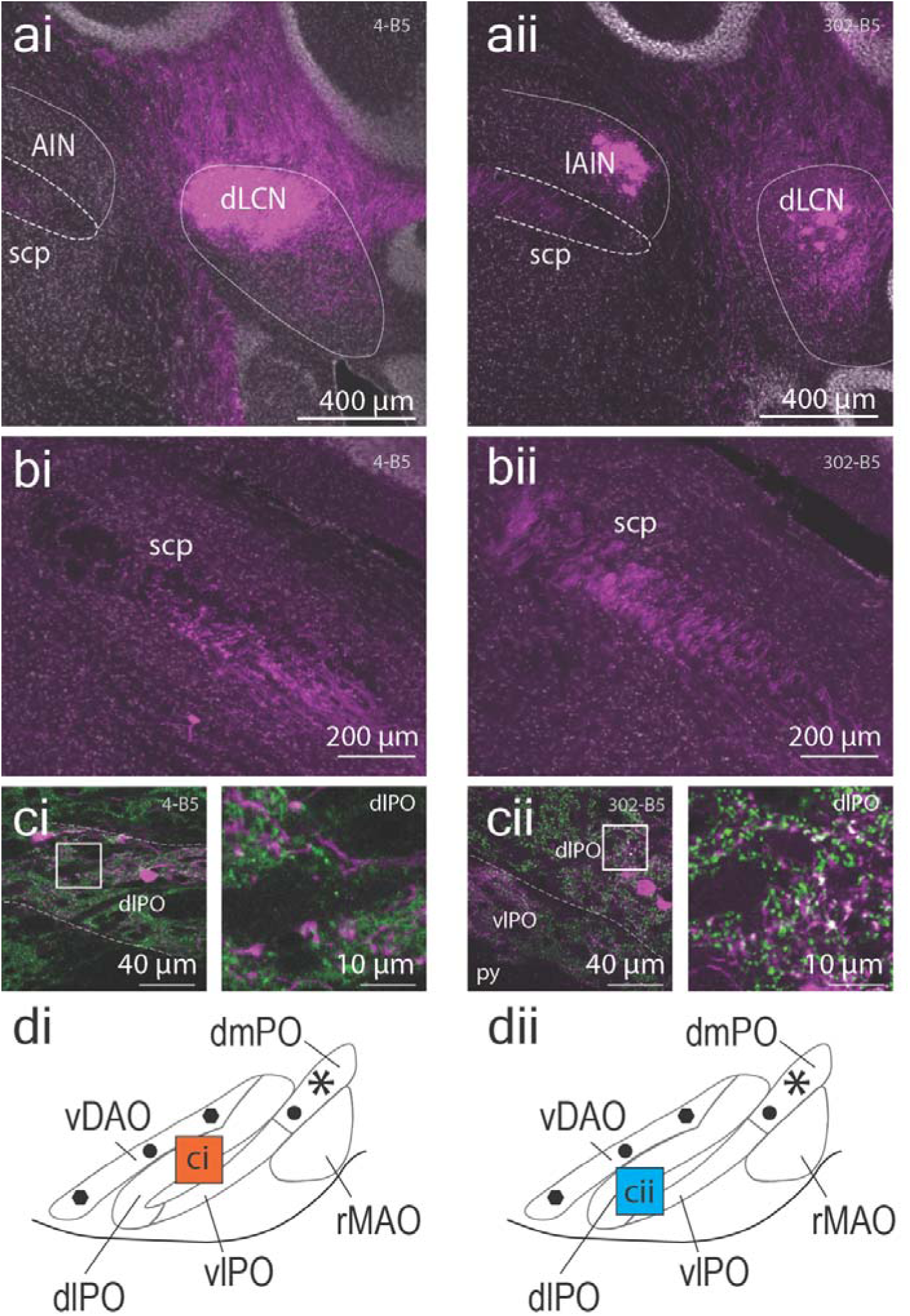
| Transsynaptic AAV1 tracing. (**a**), Example images of transsynaptic labeling in the cerebellar nuclei (CN) for injections 4-B5 and 302-B5. (**b**), Transsynaptically labeled axons in the superior cerebellar peduncle. Note the different density of labeling in bii compared to bi as well as the difference in mediolateral localization. (**c**), Images of retrogradely labelled cells in the inferior olive (IO) in the absence (ci) or presence (cii) of transsynaptically labeled GAD65/67+ varicosities of CN-IO projecting neurons. The panels on the right side of Ci and Cii correspond to the white squares indicated in the left panels. (**d**), Schematic drawing of the subnuclei within the IO, including their relation to the olivocerebellar modules as described in Figure 2e. The precise location of the confocal scans in panel c are depicted by the squares in d. The absence or presence of GAD65/67+ varicosities in IO is depicted by the color of the square, where the orange square represents an absence of feedback (segregated open-loop organization) and the cyan square represents the presence of feedback (closed-loop organization).

To find out whether there are trends in the open-versus closed-loop organization of the different modules and lobules, we next replotted all injection delineations (n = 15) on a schematic representation of the cerebellar cortex and color-coded the injections based on the level of overlap between anterograde transsynaptic and retrograde labeling we found in the IO (**Fig. 4**). When considering our injections, the D1, D2, C1, C2, C3, A2 and X-CX modules in crus 2 appear to have particularly prominent open-loop microcomplexes, whereas the D0 module in crus 2 may only have closed-loop microcomplexes. Instead, all microcomplexes of all modules in crus 1 appear to be closed-loop (**Fig. 4**). Likewise, the majority of the microcomplexes of the modules in the paramedian lobule were probably also closed-loop, while those of lobule simplex were more mixed (for details, **Table S1**).

**Fig. 4.**
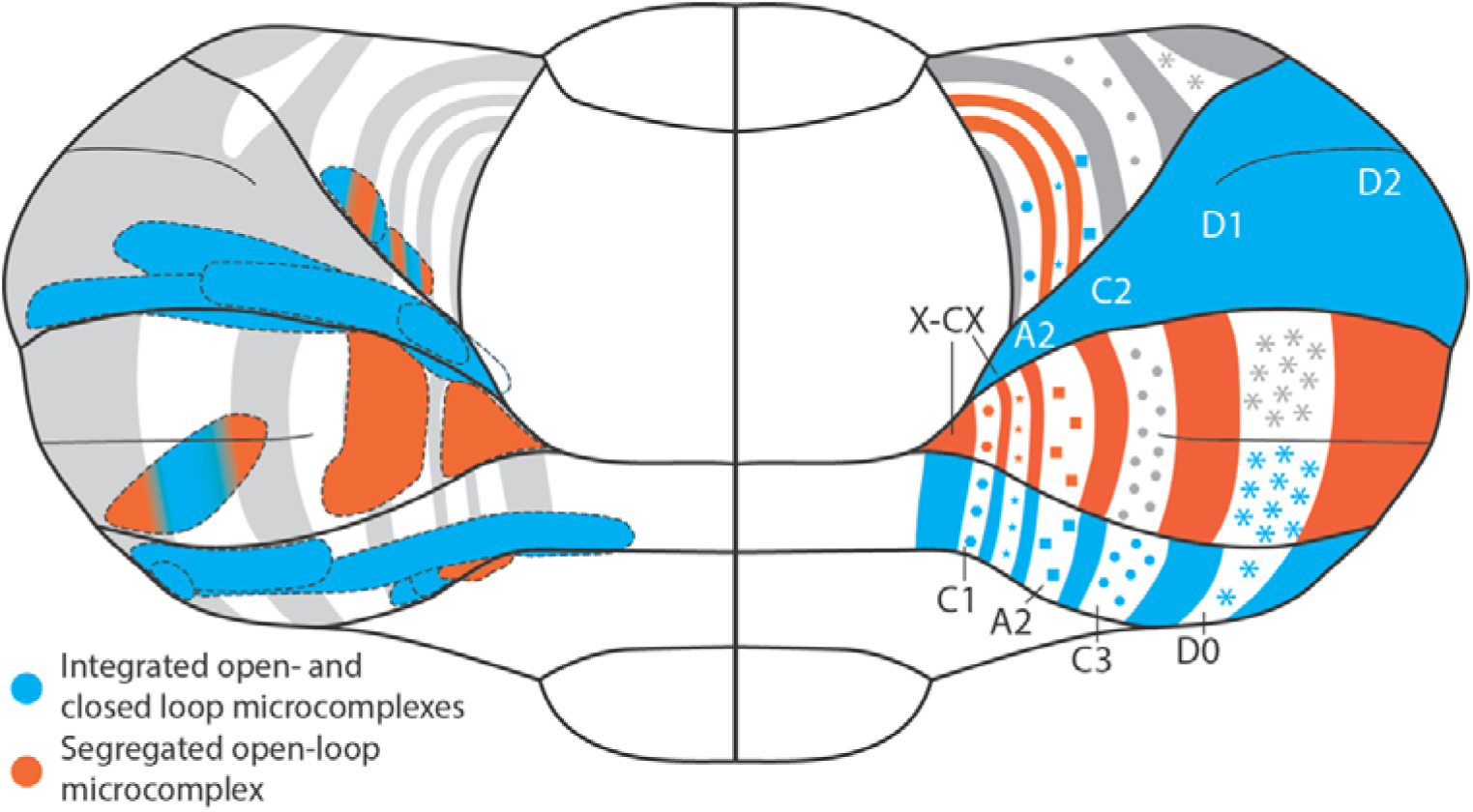
| Summary of findings regarding the existence of open-loop microcomplexes (orange) that are segregated from the typical organization (blue) where open– and closed-loop microcomplexes are fully integrated within modules of neocerebellum. On the left, the individual injection spots are colored in blue and orange when we did or did not find a significant feedback projection to inferior olive, respectively. On the right, these findings are extrapolated to visualize the seemingly lobular organization of the microcomplexes. Within our samples, segregated open-loop microcomplexes were more frequently observed in crus 2, while crus 1 and the paramedian lobule yielded predominantly closed-loop configurations. For details of names of zones, see references (Voogd *et al*. 2003, Apps and Garwicz 2005, Apps *et al*. 2018).

Our data raise the possibility that there may be differences among the different lobules in that some may be predominantly integrated in the mixed or closed-loop microcomplexes of the modules, whereas others may be more prominently engaged in open-loop microcomplexes. Even so, given the limited number of injections per module and the potential technical caveats of transsynaptic tracing, the lobular trends observed in the present study should be considered as hypothesis-generating rather than established differences in modular organization. Future studies with denser regional coverage and a comprehensive approach for the entire neocerebellum should elucidate to what extent our findings can be generalized.

To investigate the potential relevance and functional implications of open-loop microcomplexes within the neocerebellum, we expanded on a cerebellar model that contains only closed-loop microcomplexes (Fernández Santoro *et al*. 2025). In this extended model, we did not construct a fully independent open-loop circuit. Instead, we added a second population of 100 Purkinje cells to the original closed-loop model, representing the open-loop microcomplex. These cells receive distinct parallel fiber input of the specific lobule involved, but they share the same olivary input as the closed-loop microcomplex, such that their climbing fiber input is entirely determined by the inhibitory CN activity of the closed-loop circuit and not by the excitatory CN activity that forms part of a segregated open loop. As in our previous work, we focus on Purkinje cell activity, as this is the main driver of CN output, and changes at the level of Purkinje cells are sufficient to capture how differences in input structure propagate through the system.

The model focuses on the resonant network dynamics that induce long-term potentiation (LTP) and long-term depression (LTD) at the parallel fiber inputs to PCs embedded in the so-called Upbound and Downbound microzones, which are characterized by increases and decreases in simple spike (SSpk) activity during the initial stages of learning (De Zeeuw 2021). To date, these forms of LTP and LTD are predominantly occurring in the temporary absence and presence of complex spikes (CSpks), respectively, with LTP being the dominant plasticity in the Upbound microzones and LTD in the Downbound microzones. The original model with exclusively closed-loop modules receives ongoing arbitrary signals representing contextual input from 5 parallel fiber bundles, which is conveyed by 100 PCs via 40 CN cells to an IO with 40 neurons (Fernández Santoro *et al*. 2025). We expanded this model by adding another 100 PCs that are part of the open-loop microcomplex (**Fig. 5a**). The open-loop microcomplex receives parallel fiber inputs that differ in terms of source and information content from the parallel fiber inputs to the closed-loop microcomplex, while the climbing fiber input from the IO is the same for both microcomplexes. Differences in PC inputs from the mossy fiber – parallel fiber pathway are modelled as a change in input decay time, with the open-loop microcomplexes having a slower decay (**Fig. 5b**). As the input to our model network comes from bundles of parallel fibers, the parallel fiber to PC synapse acts as a “meta-synapse”, representing the average direction of change of the individual synaptic weights in the parallel fiber bundle. To capture the essential interactions between LTP and LTD we model the parallel fiber to PC synapse with a mixture of a Bienenstock-Cooper-Munro-type (BCM) learning rule and a CSpk-dependent plasticity mechanism that depends on the parallel fiber input. When we activated the PCs through their parallel fiber input, we found that in the closed-loop microcomplex CSpks only occurred when favorable oscillatory patterns emerged in the parallel fiber input. The CSpks were elicited as a resetting mechanism when there was inhibition of the IO following disinhibition of the CN (**Fig. 5c**, left). Indeed, when the parallel fiber postsynaptic currents are increased in the closed-loop microcomplexes, CN neurons of this module are disinhibited following an initial inhibition process due to increased SSpk firing, which in turn elicits climbing fiber activation in the IO, i.e., a CSpk in the PC of the corresponding microzone (Loyola *et al*. 2023). However, the relationship between the postsynaptic currents and the CSpks was not present in the open-loop microcomplexes (**Fig. 5c**, right). In fact, as the open-loop CSpks were elicited without the existence of the oscillatory parallel fiber patterns, we can conclude that these CSpks occurred due to the dynamics of the closed-loop microcomplex. This implies an organization where the open-loop microcomplexes elicit CSpks and adapt the weight of their parallel fiber to PC synapses that are based on computations done by the closed-loop microcomplex, serving as the root cause of the CSpk pattern. To mimic the characteristic mechanisms and properties of closed-loop microcomplexes of the Upbound and Downbound micromodules, we assessed oscillations of the postsynaptic currents of the parallel fiber activity before the occurrence of a CSpk using two-tailed Pearson correlations between the currents and corresponding sine waves (**Fig. 5d**). With this approach the average SSpk frequency of the Upbound and Downbound microzones were under baseline conditions set at ∼60 Hz and ∼90 Hz, respectively (**Fig. 5e, top**), which is in line with experimental findings (Zhou *et al*. 2014, De Zeeuw 2021). Moreover, upon parallel fiber activation the PCs could adjust their SSpk firing rate in a bidirectional fashion in that the average frequency increased in the Upbound microzones, whereas it decreased in the Downbound microzones (**Fig. 5e, bottom**), as predicted by theoretical considerations (De Zeeuw 2021). Likewise, the changes in the firing patterns of CN neurons (**Fig. 5f**) and IO neurons (**Fig. 5g**) upon alterations in parallel fiber activity were also both in line with experimental observations (Zhou *et al*. 2014, De Zeeuw 2021).

**Fig. 5.**
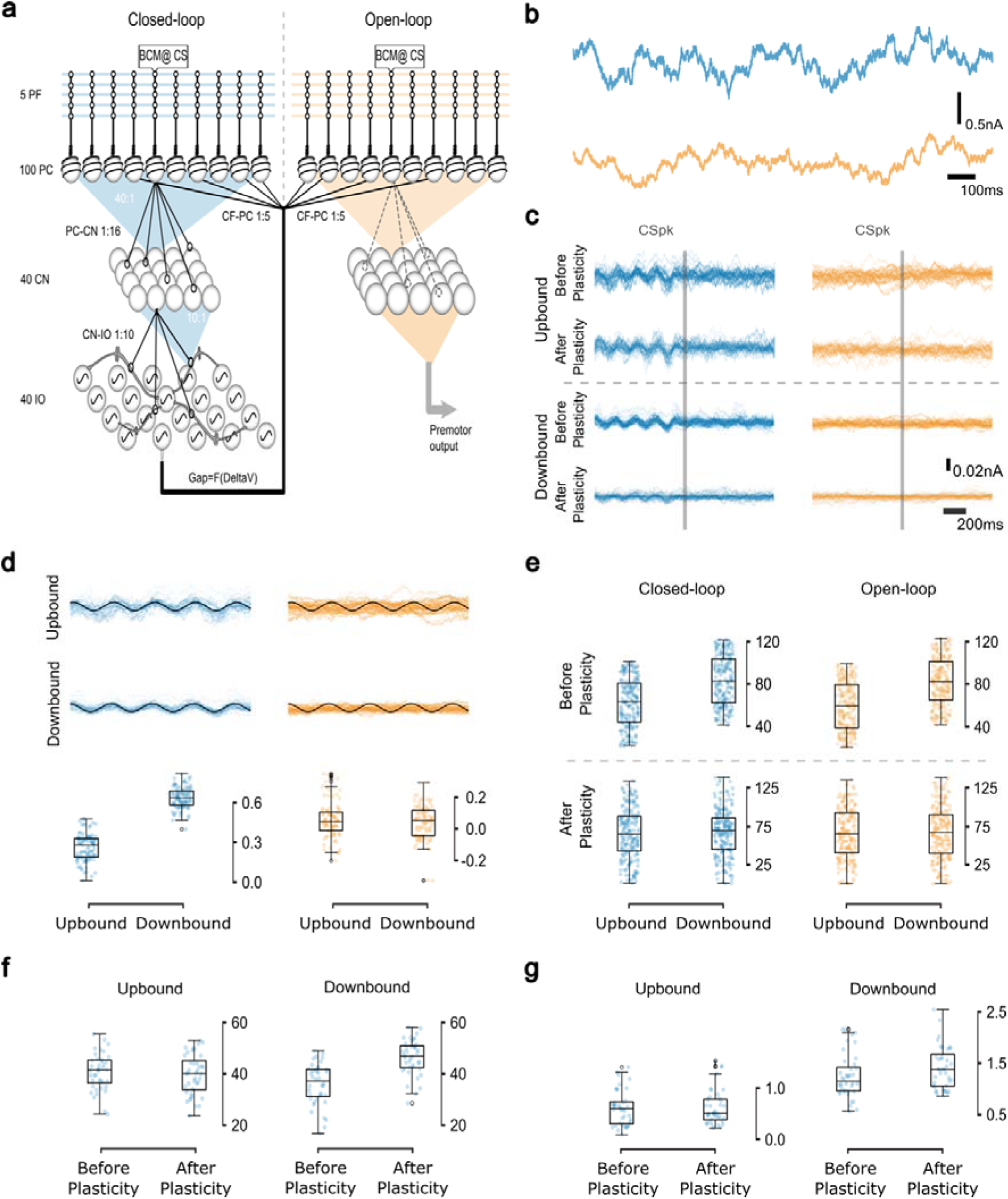
| Novel configuration allows for segregated mossy fiber – parallel fiber control of closed-loop and open-loop microcomplexes. (**a**), Diagram of the olivocerebellar loop model, which comprises 5 parallel fiber (PF) bundle inputs onto 100 PCs for both the closed-loop and open-loop microcomplex (200 PCs in total). In the closed-loop microcomplex each PC projects to 16 CNs that provide feedback to the IO and in the open-loop microcomplex each PC projects to 16 CNs that project to premotor areas. The closed-loop CN layer consists of 40 inhibitory neurons, which receive input on average from 40 PCs (from 30 to 52 PCs). Each CN neuron engaged in a closed-loop microcomplex projects to 10 IO cells. There are 40 IO cells, each of them receiving on average 10 CNs (from 6 to 16 CNs). The IO cells are modelled with gap junctions and subthreshold oscillations (Negrello *et al*. 2019). Half of the IO cells project to the PCs with each of those IO cells projecting to an average of 5 PCs from each microcomplex (from 2 to 9 cells), while each PC only receives signals from 1 IO neuron, completing the three-element micromodule. Each IO neuron connects to all other IO neurons in the population via the gap junctions. (**b**), The open-loop microcomplex (bottom) receives a mossy fiber – parallel fiber input that has a slower decay time than that of the closed-loop microcomplex (top), in line with the Ornstein-Uhlenbeck (OU) process. (**c**), In the closed-loop microcomplex a CSpk is elicited following an increase of PF-driven postsynaptic currents in the PC, which evokes a sequence of inhibition and disinhibition in the CN neurons that inhibit the IO cells. In the open-loop microcomplex the occurrence of a CSpk is purely due to intrinsic processing in the IO. This difference holds true for both Upbound and Downbound PCs, both before and after plasticity induction. (**d**), Sine waves were found to approximate the oscillations of the PF postsynaptic currents before the CSpk in the closed-loop microcomplexes for both the Upbound and Downbound microzones (top panel). As a sine wave approximation for the inputs to Upbound PCs, we used 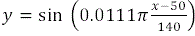, and for Downbound PCs we used 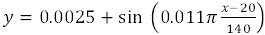 These sine waves were extrapolated to the open-loop microcomplexes. A two-tailed Pearson correlation was done between each PF postsynaptic current and its corresponding sine wave for both types of microzones and both types of microcomplexes (bottom panel). (**e**), Boxplots of SSpk frequencies of the Upbound and Downbound PCs for both the closed-loop and open-loop microcomplexes; note that the frequency in the Upbound microzone is lower than that in the Downbound microzone, while there are no significant differences between the two types of microcomplexes. (**f**), Boxplots of CN spikes before and after plasticity. (**g**), Boxplots of CSpk frequency before and after plasticity are consistent with experimental results.

To relate the plasticity dynamics of open– and closed-loop microcomplexes to a behaviorally relevant stimulus structure, we simulated delay eyeblink conditioning (Dijkhuizen *et al*. 2024) using the identical paradigm previously validated in our closed-loop model (Fernández Santoro *et al*. 2025). The paradigm is designed to reveal small plasticity-driven reconfigurations of the network steady state rather than to reproduce trial-by-trial behavioral acquisition across the entire conditioning process (see Methods). Our original simulations already demonstrated that the stable Upbound and Downbound baseline states under stationary Ornstein-Uhlenbeck (OU) inputs persist during this classical paradigm of learning-dependent timing. In our experiments a conditioned stimulus (CS) that lasted 250 ms was paired with an unconditioned stimulus (US) that co-terminated with the CS for the final 30 ms of each trial **(Fig. 6a, b**). The CS was provided via the mossy fiber – parallel fiber pathway to the closed-loop microcomplex only, while the US signals were also provided by the climbing fibers originating from the IO to both the closed-loop and open-loop microcomplex (**Fig. 6a**). Moreover, for these simulation experiments we also added the inhibition provided by the molecular layer interneurons, which is known to have an important contribution to the SSpk dynamics during eyeblink conditioning (ten Brinke *et al*. 2015, Boele *et al*. 2018). To simulate the inhibitory effect of molecular layer interneurons onto PCs (Heiney *et al*. 2014, ten Brinke *et al*. 2015), we modeled the CS as a short pulse followed by a prolonged inhibition (**Fig. 6b**; see also Methods for details). This simplification reflects the known feedforward dynamics of molecular layer interneuron inhibition that follows parallel fiber excitation (Wulff *et al*. 2009, Badura *et al*. 2013, ten Brinke *et al*. 2015). We applied the CS and US *in-silico* with similar durations, sequences and intertrial intervals as provided in physiological eyeblink conditioning experiments (Heiney *et al*. 2014, ten Brinke *et al*. 2015). Each *in-silico* conditioning block consisted of an initial US followed by 10 CS+US co-terminating pairings and a final CS with a duration of 220 ms (**Fig. 6c**), and each US reliably triggered a CSpk, setting a semi-rhythmic activity pattern (**Fig. 6d**). In line with the reported dynamics in SSpk firing after conditioning (Heiney *et al*. 2014, ten Brinke *et al*. 2015), we observed the SSpks to decrease and increase after plasticity induction in the Downbound and Upbound microzones, respectively **(Fig. 6e,f**). This suggests that the molecular layer interneuron inhibition inserted in the microzones is predominantly engaged in the Downbound PCs, while additional homeostatic mechanisms such as changes in intrinsic excitability may be implicated in the Upbound PCs in the same context (De Zeeuw 2021).

**Fig. 6.**
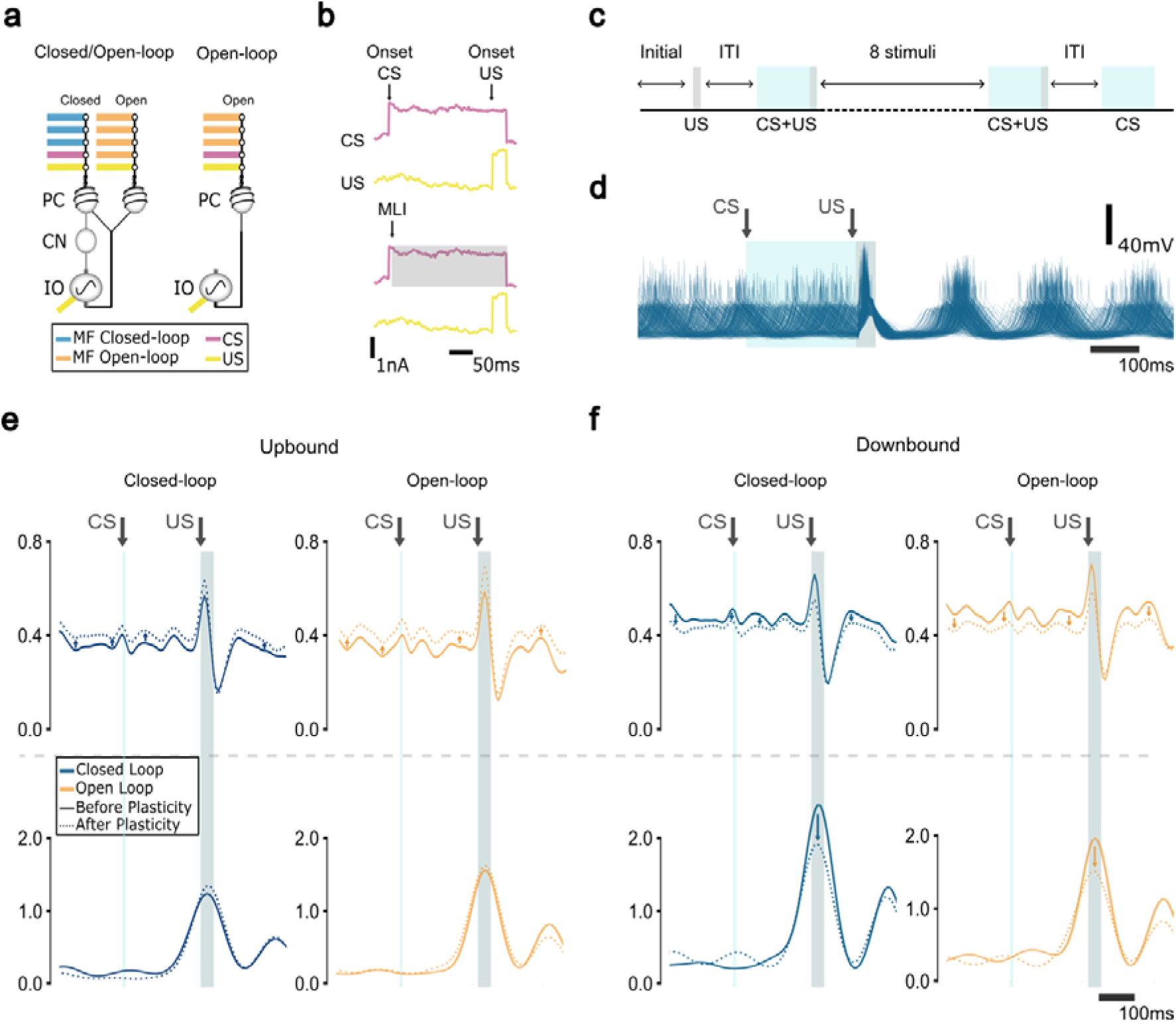
| Cerebellar conditioning in-silico. (**a**), Diagram of two different configurations of micromodules side by side, including a mixed closed– and open-loop microcomplex (left) as well as a pure open-loop microcomplex (right). It is indicated in colors how the conditioned stimulus (CS; purple) and unconditioned stimulus (US; yellow) are relayed to the different microcomplexes. In the simulations under investigation the CS enters the network only via the mossy fiber – parallel fiber pathway and the US enters the network via both the mossy fiber – parallel fiber pathway and the climbing fiber pathway. The mossy fiber – parallel fiber pathway and the climbing fiber pathway are responsible for modulating the simple spike (SSpks) and complex spike (CSpks) responses of the Purkinje cells (PCs), respectively. (**b**), Stimulation protocol of the classical delay conditioning experiment (top), in which the traces show the input current amplitude. For this experiment we modelled the effect of inhibition provided by the molecular layer interneurons (MLIs; bottom); the CS is reduced to a 3 ms pulse to simulate subsequent prolonged MLI-mediated inhibition of PCs (grey beam). (**c**), Each training block consists of one US-only trial, followed by ten CS-US paired trials with varying intertrial intervals (ITIs), and ending with a CS-only trial. (**d**), Membrane voltage traces from IO neurons in a Downbound module show robust US-evoked CSpks in the closed-loop configuration. (**e**), CS-triggered spike frequency density plots of SSpk (top) and CSpk activity (bottom) in Upbound modules. In both the closed-loop (left) and open-loop (right) configurations, SSpk firing increases after plasticity. For the CSpks, a small US-associated increase is seen in the closed-loop after plasticity, probably associated to the IO disinhibition as result of strong PC excitation. This increase is absent in the open-loop microcomplex, where both CS and US converge on the same PC population, limiting the specificity of plasticity. (**f**), Same as panel (e), but for the Downbound PCs. Here, the SSpk activity is decreased after plasticity in both the open– and closed-loop configurations. CSpk activity also decreases after plasticity in both configurations (see arrows), consistent with learning-dependent modulation of IO output. For both panels (e) and (f), we generated the CS-triggered frequency density plots using a kernel density estimate (KDE) across combined conditions (Upbound/Downbound, before/after plasticity) in a single Seaborn plot. As a result, the y-axis reflects a shared density scale across all subgroups, whereas plotting each condition separately would yield higher values due to normalization within a smaller subset.

To better understand the influence of the closed-loop microcomplex on the open-loop microcomplex we also did the same experiment without the closed-loop, where the open-loop PCs received both the CS and US through the mossy fiber – parallel fiber pathway, as well as the US signals mediated by the climbing fibers from spontaneously spiking IO neurons. Under these conditions, the relative trends of the SSpk responses resembled those seen in the closed-loop: Upbound PCs increased their SSpk activity after induction of plasticity, whereas Downbound PCs exhibited decreased firing (**Fig. 6e**, top panels). However, a notable difference emerged in the CSpk responses (**Fig. 6e**, bottom panels). In the open-loop configuration, the US did not trigger a change in CSpk activity following plasticity in the Upbound module, likely due to the lack of stimulus-specific segregation, as both the CS and US were delivered to the same PC population. Instead for the Downbound module, we observed no significant differences between open– and closed-loop configurations with relatively similar decreases in CSpk activity upon induction of plasticity (**Fig. 6f**), suggesting that intrinsic PC mechanisms are sufficient to drive the direction of plasticity.

These two configurations also have important implications for motor control. In the combined closed– and open-loop architecture, excitatory CN neurons receive converging inputs from both feedback-tuned closed-loop and open-loop PCs (**Fig. 1**). Since both sets of PCs are modulated by the CF input from the IO from the closed-loop circuit, the CN output integrates predictive error correction signals with ongoing contextual modulation. This shared IO-mediated feedback enables a coordinated, adaptive control signal that balances internal learning with flexible task demands. In contrast, in the open-loop-only configuration, the CN neurons receive input solely from the open-loop PCs. This results in a purely feedforward signal, potentially enabling faster motor responses that are less reliant on online error correction. Such a separation could support specialized control strategies, where some microzones regulate stable, learned behaviors via closed-loop feedback, and others deliver rapid context-driven responses through open-loop processing.

These comparisons demonstrate that modular segregation into open– and closed-loop microcomplexes allows the olivocerebellar system to preserve key learning dynamics even in the absence of direct feedback loops. The ability of open-loop modules to replicate the direction of SSpk adaptation while bypassing closed-loop feedback highlights a form of computational redundancy that may provide robustness under variable input configurations. Moreover, this arrangement may allow different microzones to specialize in either error correction or forward modulation depending on the context provided by the mossy fiber system, thus expanding the flexibility of cerebellar learning. Indeed, integrating open-loop microcomplexes alongside traditional closed-loop microcomplexes enriches the computational repertoire of the olivocerebellar system. In this hybrid architecture, open-loop microcomplexes can flexibly adapt their plasticity rules based on global error signals computed elsewhere in the circuit, effectively allowing peripheral open-loop microcomplexes to benefit from core closed-loop computations without directly participating in feedback inhibition. This decoupling of local plasticity from local feedback enables distributed learning across modules, supports context-dependent specialization and may increase cerebellar capacity to generalize learned associations across behavioral states. By supporting parallel learning strategies within a shared behavioral context, the mixed architecture helps balance the demands of immediate motor output and longer-term prediction.

## Discussion

Here, we provide neuro-anatomical evidence for the existence of open-loop microcomplexes within modules of the neocerebellum, next to their closed-loop microcomplexes. We first confirmed that the trajectory for axons of CN neurons that project to the IO in mice diverges from that of CN axons that project to premotor regions, with both showing different localizations in the superior cerebellar peduncle. We next showed that all AAV injections into microzones of PCs always result in retrogradely labeled cells in the IO, but not always in transsynaptically labeled varicosities in the olivary neuropil. Dependent on the lobule that is injected in the neocerebellum, there may be different parts of the superior cerebellar peduncle labeled and anterograde, transsynaptic labeling in the olivary subnuclei can be present or absent. These data indicate that the classical doctrine that modules of the neocerebellum are always completely closed might not be correct. The existence of open-loop microcomplexes segregated from closed-loop microcomplexes within individual modules raises the possibility that feedforward control of behavior and feedback control of the olivocerebellar system can be differentially coordinated. Our model of the neocerebellar system highlights that differential control of such feedforward and feedback mechanisms can enrich the computational options contributing to a behavior like Pavlovian associative conditioning.

The different trends we found in the closed-loop and open-loop configurations among the different lobules might have been partially biased by our transsynaptic approach. The use of adeno-associated viral vector AAV1-CMV-HI-eGFP-Cre-WPRE-SV40 in Ai14 reporter mice might have led to differential efficiency of labeling of the inhibitory or excitatory nature of the synaptic connections involved as well as of the anterograde vs retrograde direction of transport (Zampieri *et al*. 2014, Zingg *et al*. 2017, 2020, 2022, Lanciego and Wouterlood 2020). Moreover, the closed-loop or open-loop nature of a microcomplex by itself might yield different propensities for under-labeling, further hampering reliable quantifications. Thus, even though we used the same approach for all lobules and microcomplexes, we cannot exclude the possibility that these caveats played out differently across the different regions investigated. Given the limited number of tracing experiments and given that we have not investigated the entire cortex of the neocerebellum, the present findings should be considered as qualitative evidence for the coexistence of the two configurations within modules of the neocerebellum, rather than as robust quantifications of their relative prevalence; rigorous prevalence calculations will require multiple tracing methods with calibrated labeling efficiencies across all neocortical targets in larger populations of mice.

Despite potential caveats of our transsynaptic approach, the current description of open-loop microcomplexes within modules of the neocerebellum is in line with similar findings that have been made in the vestibulocerebellum and spinocerebellum. For example, using classical tracing approaches we have shown that some PCs in the flocculus target cells in the complex of vestibular and CN that provide feedback to the IO, whereas others only project to the vestibular nuclei neurons that drive the eye movements (De Zeeuw, Gerrits, *et al*. 1994, De Zeeuw, Wylie, *et al*. 1994). It should be noted that in the phylogenetically older vestibulocerebellum it was relatively easy to interpret classical tracing experiments, because its areas providing GABAergic feedback to the olive — e.g., the projections from the nucleus prepositus hypoglossi, ventral dentate nucleus and group y to the dorsal cap and ventrolateral outgrowth — are spatially separated from those with the excitatory neurons projecting to the oculomotor nuclei (De Zeeuw, Wylie, *et al*. 1994). Thus, unlike the neocerebellum, where these inhibitory and excitatory cells are largely intermingled in the same areas of the DCN, segregation of closed-loop and open-loop circuities can be readily analysed with conventional means in the vestibulocerebellum. Likewise, cells in the medial cerebellar nucleus, which is embedded in the spinocerebellum, can project to the cdMAO either directly or indirectly via the superior colliculus also in both an open– and closed-loop fashion (van Hoogstraten *et al*. 2024), possibly allowing for segregated feedforward and feedback control (Ruigrok and Voogd 1990, Wang *et al*. 2023, van Hoogstraten *et al*. 2024). Thus, open-loop microcomplexes within cerebellar modules may not only hold for the neocerebellum, but possibly also for older parts of the cerebellum, alluding to a function that may be well conserved through evolution. Indeed, the current finding of segregated control over the open– and closed-loops by different PCs within the same module of the neocerebellum offers intriguing possibilities for new computational operations in the olivocerebellar network responsible for higher motor and cognitive functions.

The proposed cerebellar architecture with both closed– and open-loop microcomplexes within single modules implies that the specific content of information contained within a module can vary dependent on the context involved. As different cerebellar lobules receive functionally distinct mossy fiber inputs conveying context-dependent information (Schmahmann 1996, Henschke and Pakan 2020, Nettekoven *et al*. 2024), specific conditions could increase the influence of the mossy fiber – parallel fiber bundles in the open-loop microcomplexes of certain lobules, while simultaneously decreasing the electrophysiological relevance of functionally distinct mossy fiber inputs to other lobules that are responsible for regulating activity of the closed-loop microcomplexes of the same module. Consequently, our suggested architecture, with mixed open– and closed-loop microcomplexes in a lobule-dependent pattern, raises the possibility that the precise type of cerebellar feedback to the IO can be separately determined for each module, depending on where the closed-loop microcomplexes are localized across the cerebellar cortex.

The apparent lobular organization of the open-loop microcomplexes in neocerebellum is supported by functional magnetic resonance (fMRI) imaging studies in humans (Guell and Schmahmann 2020, Nettekoven *et al*. 2024), which highlight that crus 1 and crus 2 show different representations when considering default-mode processing or attentional/executive processing. Likewise, the differential CSpk responses that can be expected based on partially segregated closed– and open-loop microcomplexes have indeed been demonstrated in the crus regions of non-human primates when subjected to a missing oddball detection task (Okada *et al*. 2025). Moreover, they might also contribute to the differential engagement of supervised and error-based learning mechanisms in tracking tasks (Streng *et al*. 2017, Raymond and Medina 2018, Shadmehr and Fakharian 2026).

Even though their climbing fiber – driven plasticity rules may be equally robust (Badura *et al*. 2013), the open-loop microcomplexes will be less influential than the closed-loop microcomplexes for shaping SSpk coding within their respective olivocerebellar module. In simplified terms, the relevance or weight of the mossy fiber – parallel fiber input to the open-loop microcomplex is to a large extent determined by the timing of the mossy fiber input to the closed-loop microcomplex, via which the CSpk pattern of the entire module is controlled. Thus, albeit the SSpks and CSpks constantly influence one another within the entire modular network, the timing of the climbing fiber activity within the closed-loop microcomplex relative to that of its mossy fiber – parallel fiber inputs may be critical for controlling the SSpk patterns of all microcomplexes (Streng *et al*. 2017). Indeed, by implementing the architecture of partially segregated open– and closed-loops into our model, the peripheral open-loop microcomplex was able to adapt the parallel fiber to PC synapses based on computations performed in the closed-loop core, thereby reducing the computational load. This internal differentiation establishes a division of labor within modules, whereby closed-loop microcomplexes refine prediction errors and stabilize feedback dynamics, while peripheral open-loop microcomplexes implement rapid feedforward adjustments based on those computations. Our simulations of the cerebellum-dependent conditioning task indicate that both open– and closed-loop configurations can acquire the CS–US association, with the architectural distinction between configurations lying in how the climbing fiber signal is shaped. In this way, open– and closed-loop specializations may allow the cerebellum to balance immediate motor execution with longer-term, predictive learning.

## Methods

### Animals

For the transsynaptic tracing experiments, we used male and female Ai14 reporter line mice (expressing TdTomato in the presence of Cre; No. 007914; Jackson Laboratories), which were maintained on a C57BL/6 background. All mice were 52–200 days old, group-housed and maintained on a 12 h: 12 h light–dark cycle with ad libitum food and water. All experimental procedures were approved a priori by an independent animal ethical committee (DEC-Consult) as required by the relevant institutional regulations of the Erasmus MC and the Dutch legislation on animal experimentation. Permission was filed under the license number AVD1010020197846.

### Stereotactic surgeries

For all intracranial injections, mice were deeply anesthetized with isoflurane (1–2%; Isoflutek, Laboratories Karizoo) and placed in a stereotaxic apparatus (Kopf Instruments). All mice received subcutaneous injections of 50 µg/kg of buprenorphine (Temgesic, Indivior) and 5 mg/kg of Rimadyl (Carprofen, Zoetis) and their eyes were protected from dehydration using DuraTears (Alcon Laboratories). The skull was exposed with a small cutaneous incision and craniotomies were drilled above the cerebellar cortex. Tracer injections were made with pulled glass capillaries (tip inner diameter: ∼8 µm; Hirschmann ring-caps). Following insertion at 500 µm/min, the capillary was left in place for 5 min before pressure injection of the tracer at 5-25 nL/min. To reduce backflow into the pipette tract, the capillary was kept in place for at least 10 min to allow for diffusion of the tracer in surrounding tissue and was then retracted at 100 µm/min. After injections, the skin was closed using Histo-acryl skin glue (B. Braun Surgical). The mice were returned to their home cage individually and monitored for at least 45–60 min while recovering on a heating pad.

### Tracers, viruses and pseudo-trisynaptic tracing

To perform the pseudo-trisynaptic tracing we injected 30 nL of AAV1-Cre (AAV1-hSyn-Cre-WPRE-hGH) aimed at lobule simplex, crus 1, crus 2, or paramedian lobule at varying mediolateral locations, with the goal to cover large parts of the lobules. Although this approach includes only one synaptic ‘jump’ (anterograde mono-transsynaptic), it provides information about three synapses, because it allows for a combined analysis of the anterograde with the retrograde labeling in a closed-loop scenario; hence the term ‘pseudo-trisynaptic tracing’. Indeed, because the anterograde transsynaptic jump, the first synapse, allows us to visualize the axons and synapses of the transsynaptically labelled cells, we can identify transsynaptic synapses (second synapse) in the disynaptic targets from the injection site, in our case inferior olive. Additionally, because of the retrograde properties of AAV1, we can identify retrogradely labelled inferior olive cells, which have a synapse in the injection site (third synapse), through which the virus has been taken up from the injection site in the cerebellar cortex.

### Immunohistochemistry

Following transcardial perfusion with 4% paraformaldehyde (PFA) 21 days after viral injection, the brains were extracted and postfixed in 4% PFA for 1.5 h at room temperature (RT). Subsequently, the brains were transferred to 10% sucrose in 0.1 M Phosphate Buffer (PB) and left overnight at 4°C, before embedding in gelatin. The brains were embedded in 14% gelatin / 10% sucrose (in milliQ), fixed in 10% formaldehyde / 30% sucrose solution for 2:30 h, and stored at 4°C in 30% sucrose (in 0.1 M PB) until being sliced on a Leica SM2000R microtome (Leica Biosystems) at 50 µm and one in four slices were taken for subsequent processing. To prepare the slices for antibody staining, they were blocked for one hour with 10% normal horse serum (NHS) and 0.5% triton in PBS at RT after rinsing in PBS. Subsequently, slices were stained for rabbit anti-KCTD12 (no. 15523-1-AP; 1:1,000, Proteintech), mouse anti-GAD67 (1:2000, No. MAB5406, Millipore), and guinea-pig anti-VGLuT2 (1:2000, No. AB2251-I, Millipore) in 2% NHS and 0.4% triton in PBS, and left 24 h at RT. Next, slices were incubated for 2 h at RT in 2% NHS and 0.4% triton in PBS containing anti-rabbit-488 secondary antibody, anti-mouse-647 secondary antibody, and anti-guinea pig-A405 secondary antibody (Alexa Fluor, Jackson ImmunoResearch Europe; 1:400 for all). Slices were then rinsed for 40 min before being mounted onto cover slides (VWR) and placed on glass slides with Mowiol (Carl Roth) or aqueous mounting media.

### Imaging and analysis

For all fluorescence experiments, the processed slices were imaged with a fluorescence Zeiss Axio slide scanner with a 10X objective. The location of injection spots was determined using the Paxinos atlas (Franklin 2001). Additionally, brain-wide projection patterns were compared to known projections of the intended target nucleus to confirm sufficient and specific targeting. For qualification of transsynaptic varicosities in IO and fibers in the superior cerebellar peduncle, we performed higher resolution 20x scans of the IO and superior cerebellar peduncle for all mice and all inferior olive– and superior cerebellar peduncle slices. We then qualified the presence or absence of retrogradely labelled IO cells, whether varicosities (transsynaptic synapses) were present in between these IO cells, and whether transsynaptically labelled CN axons were identifiable in the superior cerebellar peduncle.

### Modelling Construction

The original model built by (Fernández Santoro *et al*. 2025) is a biophysically constrained network representation of the olivocerebellar loop, comprising PCs, CNs and IO neurons. PCs and CN neurons are implemented as adaptive exponential integrate and fire (AdEX) model (Brette and Gerstner 2005); IO neurons are modelled with subthreshold oscillations and are electrotonically coupled via gap junctions, allowing them to reproduce the resonant dynamics observed experimentally. Parallel fiber input is modelled not as individual synapses but as bundles of correlated activity, simulated as Ornstein-Uhlenbeck (OU) processes to represent the temporally correlated, stochastic input that PCs receive during awake behavior. The parallel fiber-to-PC synapse undergoes plasticity governed by the combination of two rules: a Bienenstock-Cooper-Munro (BCM) rule that potentiates or depresses the synapse as a function of recent PC activity relative to a sliding threshold, and a CSpk-triggered plasticity proportional to the instantaneous parallel fiber activity at the time of each CSpk. Simulations use a “frozen-input” design, in which a single stochastic OU trace is generated once and then replayed across successive epochs — plasticity-off, plasticity-on, and plasticity-off-again — so that any difference between epochs can be attributed unambiguously to plasticity-driven reconfiguration of the network steady state rather than to differences in the input. Intrinsic excitability parameters for PCs are tuned to match the baseline SSpk firing rates reported experimentally for Upbound (∼60 Hz) and Downbound (∼90 Hz) microzones. A full description of the original model, including all equations and parameters, is given in Fernández Santoro et al. (2025); for the current study most of the parameters for the different populations are as described for the original model (Fernández Santoro *et al*. 2025). Only the numbers of parallel fibers (PFs) and PCs have changed from 5 to 10 PFs and 100 to 200 PCs, while we also implemented the molecular layer interneurons (**Fig. 6b**).

### General simulations

For all simulations the Euler-Maruyama method was used to simulate the frozen noise. For the rest of the neuron groups, the forward Euler method was used to integrate the differential equations. Two integration timesteps were used, one for the simulations (0.025 ms) and one for the recording of the neuron output (1 ms). All experiments had a simulation length of 120 s. We started by generating a seed, which included a network with connections, as well as related cell parameters and frozen PF input. The open-loop PF input had a time delay that was bigger than that in the closed-loop microcomplex.

The frozen PF input was used for the “no plasticity” experiment. For the subsequent “plasticity” epoch, we enabled the BCM and LTD/LTP mechanisms and applied the same frozen PF input. At the end of this learning phase, the evolving PF-PC synaptic weights were smoothed using a Savitzky–Golay filter (window size = 100 samples, polynomial order = 2) to reduce high-frequency noise. We then extracted the final value of each filtered weight and used it as the static weight for the next “after plasticity” simulation with the same frozen input.

### Parallel fiber bundles

The increase in the number of PFs was done to have two different sets of PF bundles, one for the closed-loop and one for the open-loop (parameter values chosen are shown in **Table 1**). We simulate the PF input, representing the activity of PF bundles, as an Ornstein-Uhlenbeck (OU) process.

**Table 1.** Parameters of the Ornstein-Uhlenbeck processes.

| Simulation | OU time constant OU (ms) |  | OU standard deviation OU |
| --- | --- | --- | --- |
| Different time constant | Closed-loop microcomplex: | 50 | 0.25 |
|  | Open-loop microcomplex: | 100 |  |

### PC neuronal model

The PC population is modeled as an adaptive exponential integrate-and-fire (AdEX) model (Brette and Gerstner 2005). All parameters are the same as for the original model, except for the intrinsic current (**Table 2**).

**Table 2.** New parameters of the PC model.

| Parameters | New | Old |
| --- | --- | --- |
| Intrinsic current $I_{int}$ (nA) | 0.85 ± 0.15[$N_{PC}$] | 0.35 ± 0.215[$N_{PC}$] |

### Connectivity

*Open– and closed-loop PF-PC*. The first 5 PFs are connected to the first 100 PCs (the open-PCs) and the last 5 PFs are connected to the last 100 PCs (closed-PCs). This is done by setting the synaptic weights to 0 when there are no synapses. Hence, the first 5 PFs only have weights >0 for the synapses with the first 100 PCs and the last 5 PFs only have weights >0 for the last 100 PCs.

*Closed-loop PC-CN connectivity*. Each closed-loop PC projects to 16 CN cells and each CN cell receives on average from 40 PCs, ranging from 30 to 52 (**Fig. 5a**). All source indices are repeated for each PC, and the CN target indices are sampled without replacement from the CN population.

*Closed-loop CN-IO connectivity*. Each CN projects to 10 IO cells and each IO cell receives on average input from 10 CN cells, ranging from 6 to 16 (**Fig. 5a**). The connectivity is generated ensuring these ratios. Cell indices are repeated for each CN cell, while IO indices are sampled without replacement from the IO population, avoiding duplicate connections.

*Open– and Closed-loop IO-PC (i.e., climbing fiber, CF) connectivity*. Each IO CF connects to an average of 5 PCs (ranging from 2 to 10) of each microcomplex (open or close) and each PC receives input from only 1 IO cell (**Fig 5a**). IO cell indices are sampled without replacement from the IO population, ensuring each PC is either uniquely assigned to a source or connected to a randomly chosen source from the preselected pool.

### Eyeblink conditioning in silico experiment

We used step currents of different amplitudes and durations (**Table 3**) to simulate the conditioned stimulus (CS) and unconditioned stimulus (US). For the CS we added a step current of 250 ms to PF 4, while for the US we added a step current of 30 ms to PF 5. The US was copied onto the IO with a 4 ms delay and multiplied by 2.8 to have the desired effect on the IO spike response. The ITI was randomly chosen between 8-12 seconds. The CS went to only the closed-loop microcomplex, while the US signals were conveyed to both microcomplexes (**Fig. 6a**). As the CS pulse increases the activity of both PCs and molecular layer interneurons (MLIs), because of their common PF input, the PC will be quickly inhibited by the MLIs. We modelled this feedforward inhibition effect by making the CS pulse last 3ms. The US pulse stayed the same and the distance between the two stayed the same (CS starts 220 ms before the US).

**Table 3.** Parameter values for the different micromodules.

| Stimulus | Pulse amplitude (nA) | Pulse duration (ms) |
| --- | --- | --- |
| CS | 1.5 | 3 |
| US | 1.5 | 30 |

### Only Open-loop microcomplex

We modelled the CS-US experiment also only on the open-loop microcomplex (without the effect of the closed-loop microcomplex). In this scenario we have open-loop PCs and the same IO as for the other experiments. Both CS and US were received by the open-loop PCs and as before a copy of the US was received by the IO. However, in this case, the synapses in the closed-loop microcomplex between PC-CN-IO were turned off so that the IO did not receive any input from the closed-loop microcomplex. As such, we kept the IO firing frequency by changing the baseline of the OU process of the 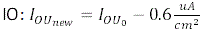.

---

**Table S1.** Quantification of labeling in the inferior olive subnuclei (20x), including the number of cells counted per IO subnucleus and an estimate of the amount of varicosities that could readily be observed. These data were used to delineate the injection sites in the schematic overviews (Figs. 2-4). The letters “a-e” refer to the codes for the slices used; numbers (e.g., “-1 or 5”) refer to the rostrocaudal level implicated, with higher numbers indicating more rostral levels. Abbreviations: ca/bMAO, subnuclei b and a of caudal medial accessory olive; ccMAO, subnucleus c of caudal medial accessory olive; cdMAO, subnucleus d of the caudal medial accessory olive; dlPO, dorsal leaf of the principal olive; dDAO, dorsal fold of the dorsal accessory olive; dmPO, dorsomedial group of the principal olive; iMAO, intermediate part of the medial accessory olive; rMAO, rostral medial accessory olive; vDAO, ventral fold of the dorsal accessory olive; vlPO, ventral leaf of the principal olive; vlo, ventrolateral outgrowth.

## Acknowledgments

We thank N. van Wingerden for performing the injections in the cerebellar cortex.

## Author Contributions

Conceptualization, WSv, EMFS, and CIdZ; methodology, WSvH and EMFS; software, WSvH and EMFS; validation, WSvH, EMFS, and CIdZ; formal analysis, WSvH and EMFS; investigation, WSvH; resources, CIdZ; data curation, WSvH and EMFS; writing—original draft preparation, WSvH; writing—review and editing, WSvH, EMFS, and CIdZ; visualization, WSvH, EMFS, and CIdZ; supervision, WSvH and CIdZ; project administration, WSvH; funding acquisition, CIdZ. All authors have read and agreed to the published version of the manuscript.

## Institutional Review Board Statement

The animal study protocol was approved by the Dutch Ethical Committee for animal experiments and performed in accordance with the Institutional Animal Care and Use Committee (Erasmus MC).

